# Dynamic balance of sparse flux vectors for efficient simulation of culture dynamics and metabolic network reduction

**DOI:** 10.64898/2026.06.17.733012

**Authors:** Ignacio Tapia García, Cristóbal Torrealba, Ricardo Luna, José Ricardo Pérez-Correa, Pedro A. Saa

## Abstract

Dynamic Flux Balance Analysis (DFBA) enables simulation of microbial culture dynamics under changing environmental conditions, but remains computationally expensive for tasks such as parameter calibration and fermentation optimization when applied using genome-scale metabolic models (GEMs). To address this challenge, we introduce Dynamic Flux Vector Balancing (DFVB), a reformulation of DFBA that solves an equivalent problem using a pre-computed, sparse basis of flux solutions that reduces the dimensionality of the internal optimization problem without information loss. Notably, DFVB provides a compact, interpretable representation of flux states that can readily identify dynamically inactive pathways and enable simulation-based automatic metabolic network reduction. We showed that DFVB produces the same culture dynamics as DFBA across multiple model scales and conditions, and identifies inactive reactions more accurately than Flux Variability Analysis (FVA) when compared to transcriptomic data profiles. Furthermore, computational performance analyses demonstrated that integrating DFVB with solver warm-start strategies and model reduction enhances computational efficiency relative to DFBA, yielding up to 3-fold reductions in simulation time for large-scale metabolic models. Finally, kinetic parameter estimation of culture dynamics with DFVB in two fermentation scenarios using a large-scale yeast GEM reached equal or higher prediction fidelity and narrower confidence intervals than DFBA, indicating improved parameter identifiability and robustness. Together, these results position DFVB as a scalable, robust, and biologically coherent framework for dynamic metabolic modeling, easing the integration of GEMs for culture dynamics simulation.

## Introduction

Genome-scale metabolic models enable the quantitative simulation of cellular physiology by integrating thousands of biochemical reactions and metabolites within a comprehensive optimization framework (Orth, Thiele, et al. 2010). These models have become indispensable tools for guiding metabolic engineering, strain design, and optimization of biotechnological processes ranging from commodity production to pharmaceuticals and enzyme manufacturing, supported by constraint-based modeling methods such as Flux Balance Analysis (FBA) (Gotsmy et al. 2024; Gu et al. 2019; Sánchez et al. 2014). The dynamic extension of this approach, Dynamic Flux Balance Analysis (DFBA), enables simulation of the temporal evolution of the cell culture under changing environmental conditions (Mahadevan, Edwards, et al. 2002). However, DFBA requires solving a large number of coupled ordinary differential equations and linear programming problems, resulting in a substantial computational burden that limits its scalability and practical integration into complex cell culture models.

Recent studies have focused on overcoming computational limitations by enhancing solver performance, introducing nonlinear reformulations through Karush–Kuhn–Tucker (KKT) conditions (de Oliveira et al. 2023), interiorpoint optimization methods (Scott et al. 2018), Runge–Kutta integration schemes (Schroeder and Saha 2020), and hybrid approaches incorporating metabolomic constraints (Dromms et al. 2020). Despite these advances, the computational efficiency of DFBA simulations remains limited to specific applications, as most approaches focus on improving numerical optimization rather than addressing the structural complexity of the underlying metabolic model (Gotsmy et al. 2024; Höffner et al. 2013).

To tackle the above difficulties, various strategies have been proposed for reducing the complexity of metabolic networks. These methods include topological reduction techniques like *redGEM* (Ataman et al. 2017), identification of topologically independent modules (Martínez et al. 2022) and reaction lumping (Erdrich et al. 2015), which systematically generate consistent *core* models by focusing on specific metabolic subsystems of interest or compressing linear pathways. Other frameworks, such as the Dynamic Reduction of Unbalanced Metabolism (DRUM) (Baroukh et al. 2014), partition the network into quasi-steady-state modules connected by dynamic metabolites. Static analysis methods like Flux Variability Analysis (FVA) are also often utilized to identify and prune blocked reactions (Mahadevan and Schilling 2003), while minimal mass-balance representations offer an alternative path for network reduction (Singh and Lercher 2020). Specifically, formulations like Alpha Spectrum (Wiback et al. 2003), MinSpan (Bordbar et al. 2014), Fast-SNP (Saa and LK Nielsen 2016) and METACONE (Altamirano et al. 2026) aim to represent the feasible flux space through extreme or sparse bases, substantially lowering the dimensionality of subsequent LP-based flux analyses while theoretically preserving biochemical accuracy. However, a common drawback of various approaches is that they frequently rely on *a priori* definitions of subsystems or make assumptions about the metabolic state, which may fail to reliably capture the transient (de)activation of pathways inherent to dynamic fermentation processes. Moreover, the robust and efficient application of these model reformulation and reduction techniques to dynamic DFBA simulations, particularly those based on minimal mass-balance descriptions, has received limited attention and remains to be evaluated.

This study addresses these limitations by introducing Dynamic Flux Vector Balancing (DFVB), a computational framework designed to structurally reduce model complexity and improve the efficiency of dynamic simulations in genome-scale metabolic models. By computing a sparse null-space basis of the flux solution space, the proposed method isolates the dynamically active metabolic subspace under a given environmental context. This reformulation not only preserves the prediction fidelity of conventional DFBA, but also enables a systematic, simulation-based reduction of the network, identifying and removing reactions that remain inactive throughout the simulated process. Furthermore, this approach enhances the identification of kinetic parameters from experimental fermentation data, leading to lower prediction uncertainty. Finally, driven by model reduction, this method is scalable to high-dimensional GEMs. The resulting framework substantially reduces computational complexity, enabling the robust integration of dynamic metabolic modeling into standard bioprocess simulation environments.

## Results

### The Dynamic Flux Vector Balancing (DFVB) Framework

Dynamic Flux Vector Balancing (DFVB) provides a rigorous mathematical reformulation of standard Dynamic Flux Balance Analysis (DFBA) designed to accelerate simulations of culture dynamics without losing precision (Fig. 1). Instead of computing the full vector of reaction fluxes (**v**) at each integration step, DFVB maps the feasible flux space onto the right null-space of the stoichiometric matrix (**N**). This basis is computed using a Rank-Revealing LU (RRLU) decomposition to preserve matrix sparsity (refer to Methods), thereby easing its biological interpretation and accelerating computational simulation. By parameterizing the flux distribution as a linear combination of null-space basis vectors, weighted by a vector of coefficients (***α***), this algebraic transformation implicitly satisfies the intracellular quasi-steady-state mass balances (Eq. 1). Effectively, this reduces the problem dimensionality from the total number of reactions (*m*) in standard FBA to (*m* − *k*) + *z* variables, where *k* represents the rank of the stoichiometric matrix, *z* denotes the number of exchange reactions, and typically *k* ≫ *z*. Consequently, the total number of constraints decreases from *n* + 2*m* to *z* + 2*m*, where *n* is the total number of metabolites. Crucially, this reformulation is not an approximation, but an exact transformation that yields an identical optimal solution to the original FBA problem while significantly reducing the complexity of the constraint set.

**Figure 1.**
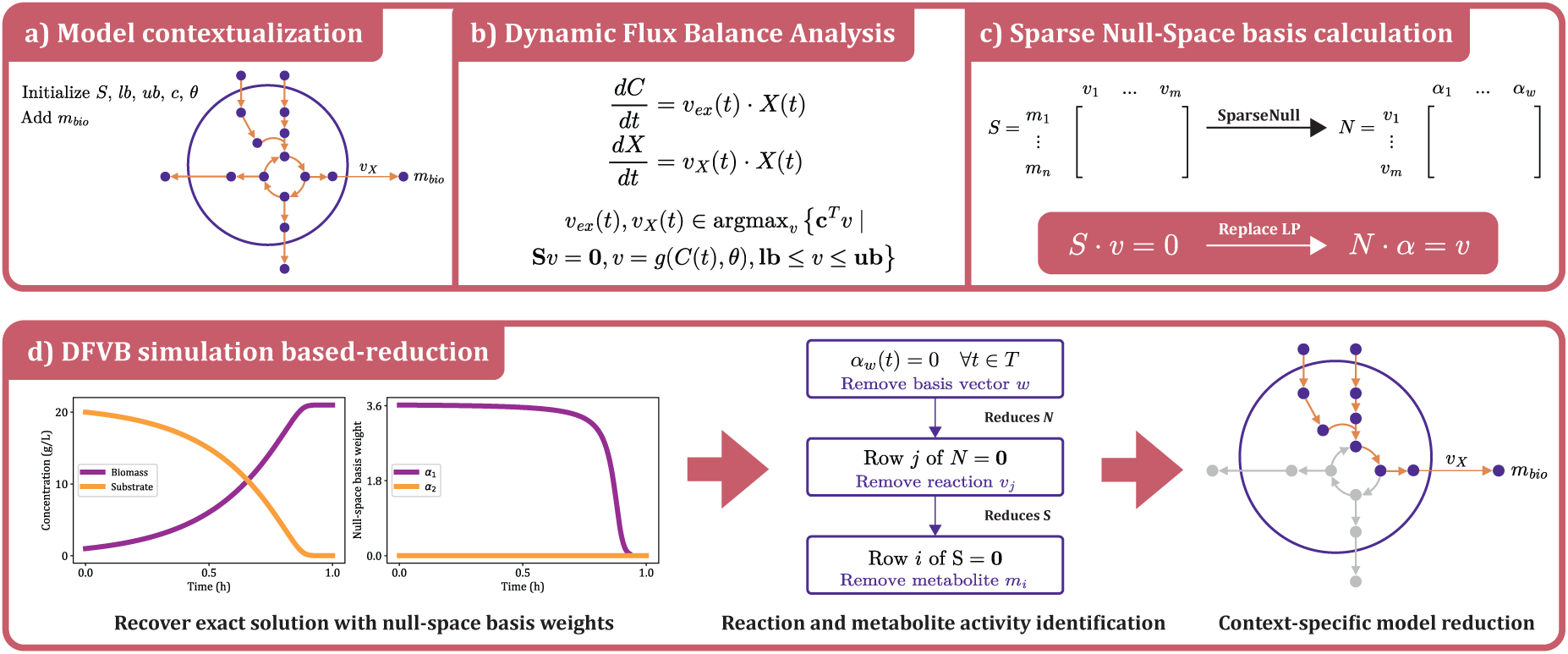
General framework for model reduction using the Dynamic Flux Vector Balancing (DFVB) methodology. (a) Contextualization and initialization of the metabolic model. (b) Standard formulation of Dynamic Flux Balance Analysis (DFBA) coupling differential equations with iterative FBA solutions. (c) Computation of a sparse null-space basis (**N**) from the stoichiometric matrix (**S**), reformulating the internal LP by replacing mass-balance constraints with a representation of fluxes (**v**) parameterized by latent variables (***α***). (d) Simulation-based model reduction using DFVB: monitoring the temporal dynamics of ***α*** enables the systematic identification and removal of inactive basis vectors, reactions, and metabolites, yielding a context-specific reduced model that captures the exact dynamic solution.

The parameterization of fluxes through latent variables provides in addition a direct mechanism for systematic, simulation-based model reduction (see Alg.1). Because each coefficient *α_j_* (*t*) reflects the activation of its corresponding flux vector basis, DFVB simulations can identify dynamically inactive pathways. Basis vectors for which *α_j_* (*t*) = **0** across the entire simulation window (*t* ∈ [0*, T*]) are classified as inactive. The sequential elimination of these inactive basis vectors from **N**, followed by the removal of the resulting zero-flux reactions and orphan metabolites from **S**, produces a highly compact, context-specific stoichiometric network. This algorithm retains all dynamically accessible biological pathways while eliminating irrelevant degrees of freedom. This significantly reduces the computational demand required for large-scale dynamic modeling, transcriptomic analysis, and kinetic parameter estimation.

### Dynamic flux balance simulations with DFVB are accurate

In order to evaluate the accuracy of DFVB relative to DFBA, simulations were performed using four metabolic models under multiple fermentation conditions. The first evaluation employed the simple illustrative model proposed by Zhao et al. (2017), shown in Fig. 2A with its corresponding null-space basis. As presented in Fig. 2B, DFVB and DFBA produced identical dynamic profiles for all simulated variables, exhibiting numerical agreement up to machine precision. The associated ***α***-dynamics revealed that only a single basis vector remained active throughout the simulation, corresponding to the flux mode responsible for biomass formation. The second basis vector, associated with product formation, remained inactive. Despite eliminating four reactions and three metabolites after removal of inactive columns from the null-space matrix, the reduced model reproduced the exact full-system trajectory. This suggests that DFVB preserves the essential dynamic behavior of systems when the optimal flux distribution is unique, enabling exact simulation without loss of information due to model reduction.

**Figure 2.**
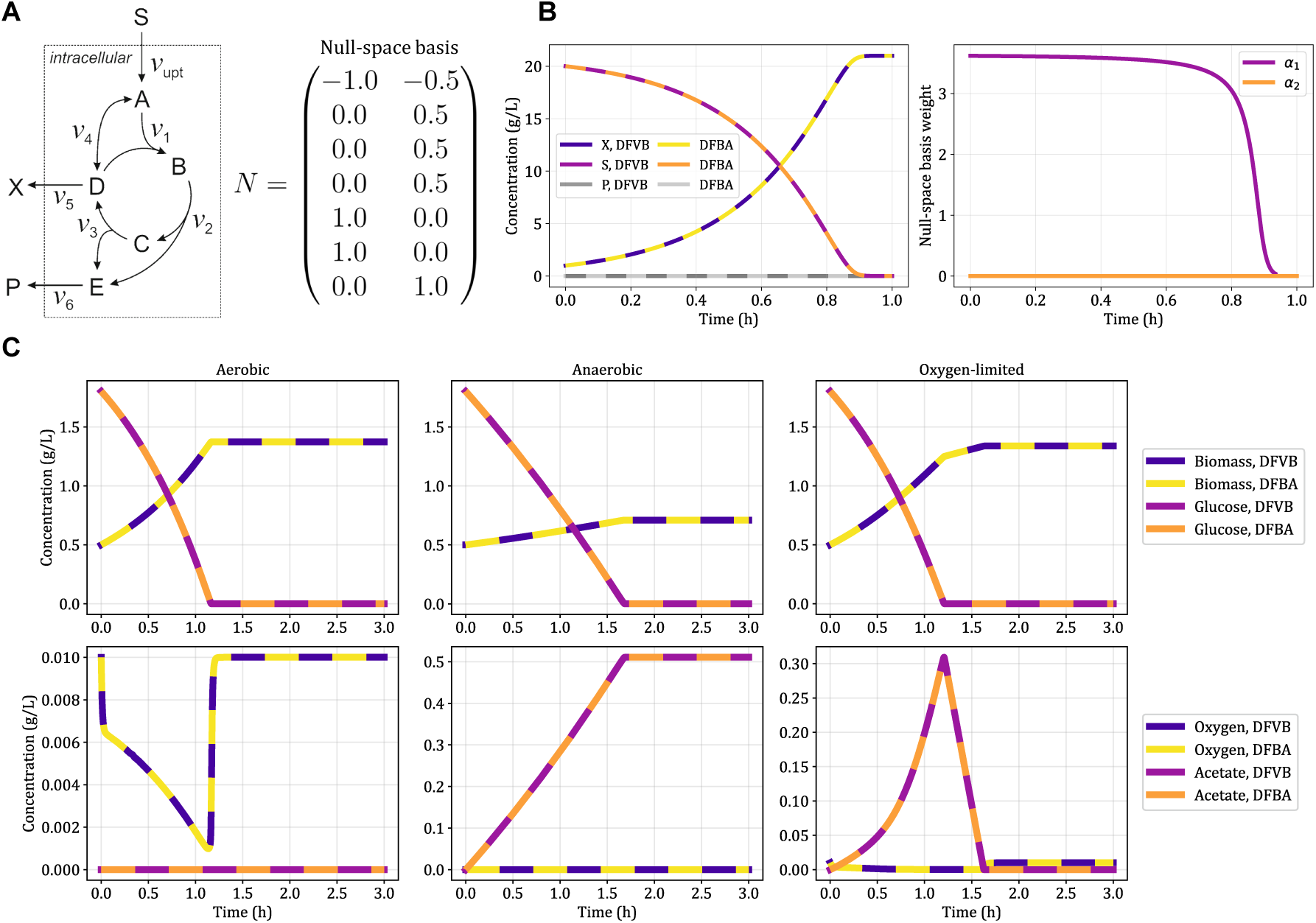
Comparison of DFBA and DFVB simulations using a simple illustrative model and the *E. coli core* metabolic network. (A) Schematic representation of the simple illustrative model, where *S*, *X* and *P* denote the substrate, biomass and a metabolic product, respectively. N represents a sparse null-basis of the stoichiometric matrix. (B) Concentration profiles of *S*, *X*, and *P* obtained with DFBA and DFVB, together with the associated dynamics of *α*. (C) Simulated concentration trajectories of biomass, glucose, oxygen, and acetate for the *E. coli core* model under aerobic, anaerobic, and oxygen-limited conditions using both DFBA and DFVB. The results of both methods are identical.

A similar analysis was performed using the *E. coli* core metabolic model under aerobic, anaerobic, and oxygen-limited regimes (Fig. 2C). In all three scenarios, DFVB and DFBA displayed perfectly overlapping trajectories for biomass, glucose, oxygen, and acetate. Under oxygen-limited conditions, both frameworks identically predicted acetate accumulation during exponential growth, followed by a metabolic shift toward acetate consumption once glucose was depleted. This diauxic behavior is characteristic of *E. coli* under glucose overflow (Pinhal et al. 2019). Table S1 in the Supplementary File summarizes the accuracy of DFVB relative to DFBA across the illustrative model, the *E. coli* core model, *i*ML1515, and Yeast9. For all models examined, DFVB reproduced the DFBA trajectories with RMSE values close to numerical precision. The variables directly constrained by the objective function, namely biomass and glucose, exhibited the smallest discrepancies across all models and conditions. In the aerobic simulations of the genome-scale models (*i*ML1515 and Yeast9), slightly larger RMSE values were observed in the production profiles of by-products (see Supplementary File Text S7), though these deviations did not substantially affect the overall simulated optimal trajectories.

### Computational Performance Evaluation

The computational efficiency of DFVB was evaluated across the *E. coli* core model, *i*ML1515, and Yeast9. Following a single DFVB simulation, each metabolic model was reduced according to the subset of active reactions and metabolites (Algorithm 1). The extent of this structural reduction is summarized in Table 1. While the reduction was modest for the *E. coli* core model, the genome-scale models showed substantial compression: approximately 14% and 28% of the basis vectors remained active for *i*ML1515 and Yeast9, respectively. Consequently, nearly 74% of reactions and 68% of metabolites were eliminated in *i*ML1515, and 68% of reactions and 64% of metabolites were removed in Yeast9.

**Table 1.** Model reduction statistics across growth conditions.

| Reduction metric | *E. coli* core |  |  | *i*ML1515 |  |  | Yeast9 |  |  |
| --- | --- | --- | --- | --- | --- | --- | --- | --- | --- |
|  | Aerobic | Anaerobic | O<sub>2</sub>-limited | Aerobic | Anaerobic | O<sub>2</sub>-limited | Aerobic | Anaerobic | O<sub>2</sub>-limited |
| Total basis vectors | 28 | 28 | 28 | 867 | 867 | 867 | 1538 | 1538 | 1538 |
| Active basis vectors | 13 | 9 | 16 | 124 | 122 | 125 | 429 | 425 | 431 |
| Reactions eliminated | 23 | 34 | 17 | 1999 | 1999 | 1997 | 2792 | 2856 | 2792 |
| Reactions conserved | 73 | 62 | 79 | 714 | 714 | 716 | 1339 | 1275 | 1339 |
| Metabolites eliminated | 8 | 13 | 5 | 1268 | 1266 | 1266 | 1832 | 1889 | 1830 |
| Metabolites conserved | 65 | 60 | 68 | 610 | 612 | 612 | 974 | 917 | 976 |

To maximize computational efficiency during dynamic integration, a sequential warm-start strategy can be used for both the standard DFBA and the proposed DFVB algorithms. Because the metabolic shifts between consecutive time steps are typically small, the optimal basis from the previous integration step is retained and supplied to the linear programming solver. However, its synergistic combination with the structural dimensionality reduction of DFVB ensures that successive optimization problems are solved with minimal pivoting. This provides a highly accelerated foundation for the dynamic analyses. Table S2 in the Supplementary File reports the CPU time metrics. Warm-starting the linear solver markedly enhanced computational performance for both approaches. For *i*ML1515 and Yeast9, disabling warm-starts increased the computational cost of DFVB by up to fivefold. Model reduction further contributed to computational efficiency; under aerobic conditions for *i*ML1515, network reduction decreased DFBA runtime approx. 3-fold (from 2.505 s to 0.807 s), and 8-fold in DFVB (from 7.594 s to 0.942 s). Similar proportional decreases were observed for Yeast9, where runtimes consistently fell to less than a third of the unreduced counterparts. Since this reduction relies on identifying the active null-space basis vectors during a preliminary DFVB simulation, DFVB emerges not merely as a simulation algorithm, but as a pre-processing step for identifying accurate and reduced model structures.

### Biologically-relevant network reduction

The functional coherence of the reduced *i*ML1515 model was evaluated using experimental transcriptomic data from Sastry et al. (2019). The subset of inactive reactions predicted by DFVB was compared against those identified by Flux Variability Analysis (FVA) using standard binary classification metrics across 49 transcriptomic conditions (Fig. 3).

**Figure 3.**
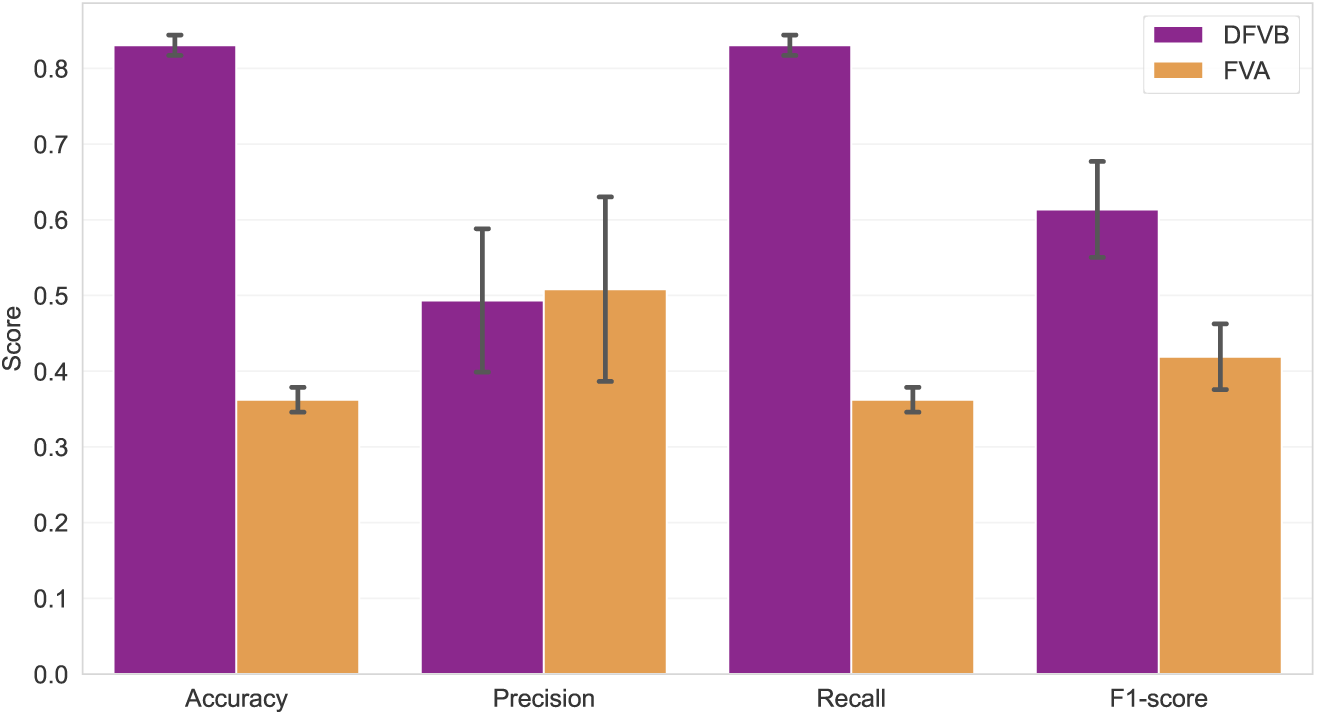
Inactive reaction classification using DFVB pre-processing and FVA.

The comparative analysis revealed that model reduction using DFVB achieved substantially higher accuracy, recall, and a superior overall F1-score compared to the FVA-based approach. However, the aggressive structural reduction applied by DFVB also resulted in a higher number of false positives (i.e., active reactions incorrectly classified as inactive). Conversely, FVA was more conservative, yielding fewer false positives but at the cost of a significantly lower recall.

### Kinetic Parameter Estimation Using Dynamic Flux Simulations

To evaluate parameter estimation capabilities, DFVB and DFBA were compared using aerobic batch fermentation data from Sánchez et al. (2014) (Fig. 4). While both frameworks captured the general dynamic trends, DFVB yielded an ethanol trajectory that more closely followed the experimental data, supported by a narrower confidence region (RMSE 0.906 vs. 1.093). DFVB also achieved a reduction in biomass prediction error (RMSE 0.086 vs. 0.128). Conversely, DFBA displayed marginally lower RMSE values for glucose, citrate, and lactate.

**Figure 4.**
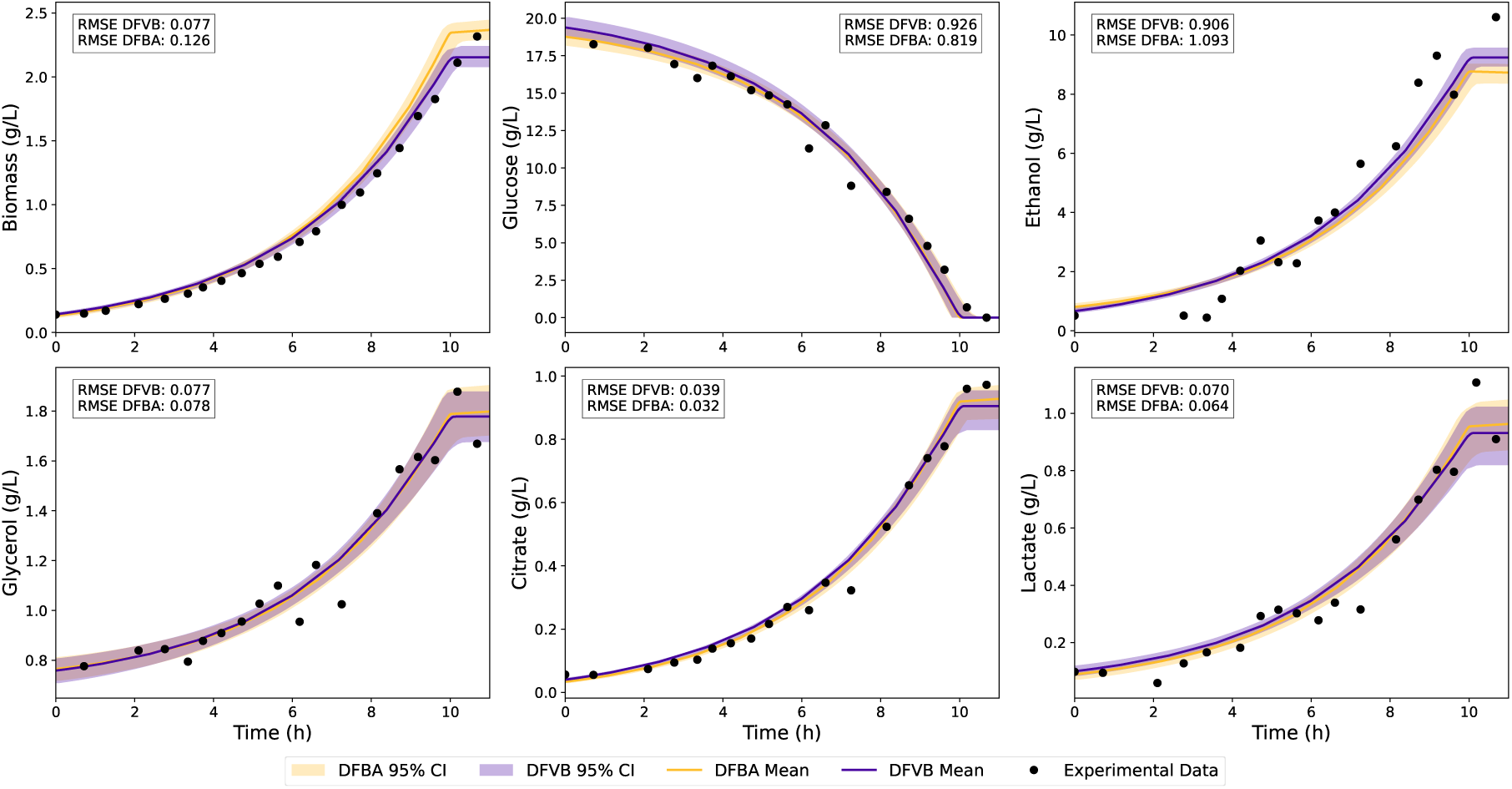
Simulation results comparing DFVB and DFBA using the experimental dataset from Sánchez et al. (2014). Shaded areas represent 95% confidence regions computed using bootstrapping, while solid lines indicate the mean prediction estimates and points denote the experimental data. RMSE values are reported for both frameworks.

Table S3 in the Supplementary File presents the parameter estimation statistics. Most initial conditions and yield coefficients converged to similar magnitudes in both frameworks. However, the lower bound for oxygen uptake 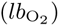 diverged substantially, with the DFBA estimate averaging 0.737 mM gDW^−1^ h^−1^ compared to 0.178 mM gDW^−1^ h^−1^ for DFVB. Additionally, both frameworks exhibited large standard deviations for the glucose saturation constant (*K_g_*) and the ethanol inhibition constant (*K_ie_*), though DFVB maintained narrower relative confidence intervals for these parameters.

A second evaluation was conducted using laboratory-scale *Sauvignon Blanc* fermentation data (Fig. 5). In this scenario, DFBA failed to accurately predict ethanol evolution, exhibiting large deviations from the experimental measurements. In contrast, DFVB achieved consistently lower RMSE values for every measured variable, including an 81.5% error reduction for ethanol, 27.8% for glucose, and 50% for primary amino nitrogen (PAN). Parameter estimation statistics (Table S4 in Supplementary File) revealed significant discrepancies in the metabolic yield coefficients between the methods. Most notably, the nitrogen yield (*Y_xn_*) was estimated at 36.14 gDW g^−1^ by DFVB compared to 14.58 gDW g^−1^ by DFBA, with DFVB reducing the relative confidence interval width by 1.8-fold.

**Figure 5.**
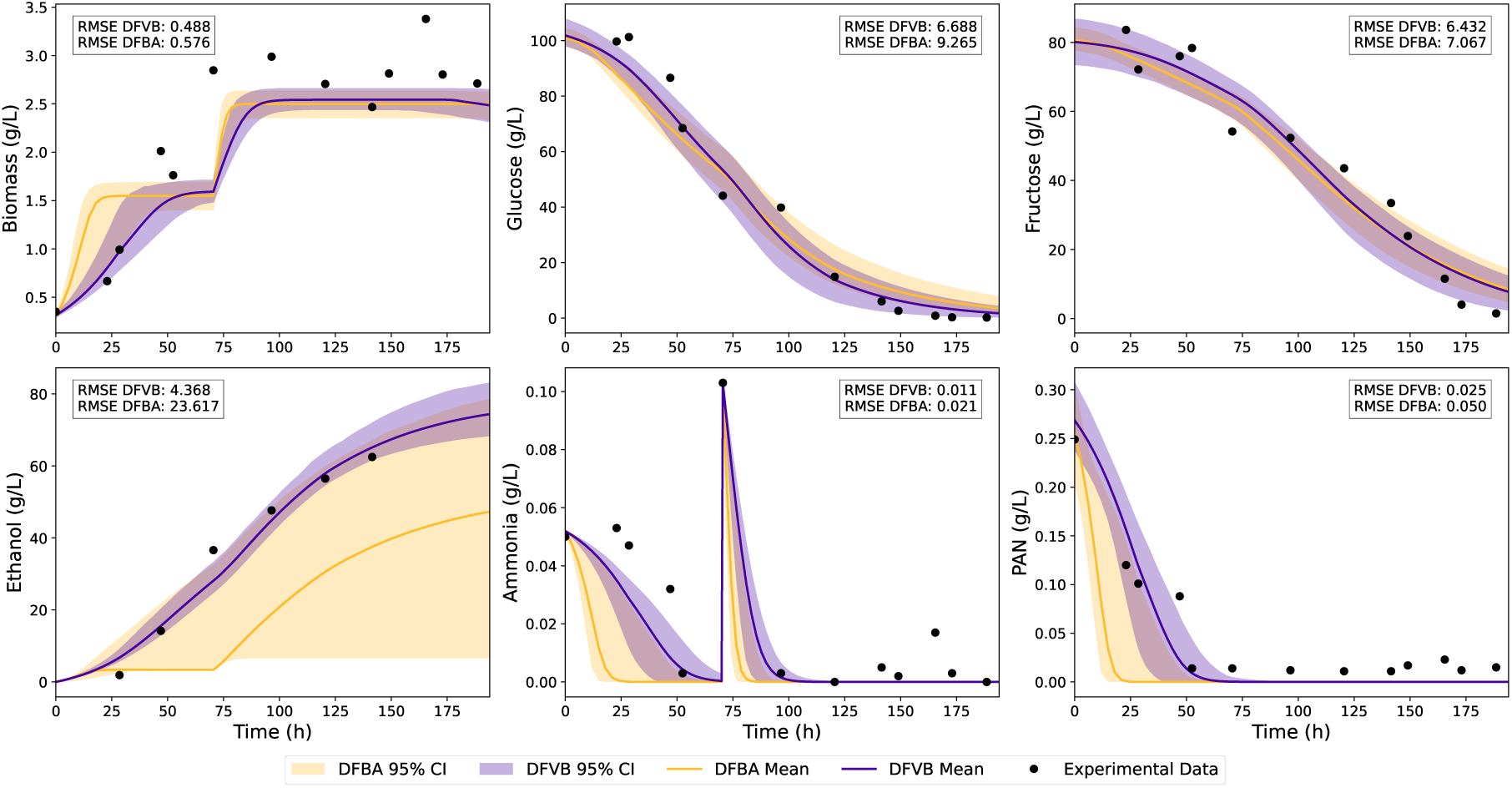
Bootstrap analysis results comparing DFVB and DFBA using the experimental dataset from Concha y Toro winery. Shaded areas represent 95% confidence regions, while solid lines indicate the mean parameter estimates and points the experimental data. RMSE values are reported for both frameworks.

## Discussion

This study introduced and evaluated DFVB, a computational framework for dynamic metabolic flux simulations designed to overcome the structural limitations of conventional approaches like DFBA. By reformulating the high-dimensional differential-algebraic system into a reduced-order model based on a sparse null-space basis, we demonstrated that DFVB retains the full simulation capability of standard DFBA while offering critical advantages in model reduction and parameter identifiability.

Our accuracy benchmarks confirmed that DFVB reproduces DFBA trajectories to machine precision when the optimal flux distribution is unique, thus preserving the exact essential dynamic behavior of the system even after model reduction. The minor discrepancies observed in the by-product profiles of genome-scale models (*i*ML1515 and Yeast9) can be attributed to the larger number of degrees of freedom inherent to these reconstructions. As noted by Smallbone and Simeonidis (2009), highly interconnected networks often possess a degenerate solution space where multiple distinct flux distributions satisfy the same optimal objective value. Under these conditions, DFBA and DFVB may naturally follow different—but equally valid—optimal internal trajectories.

From a computational perspective, the initial burden of computing the null-space projection was effectively mitigated by coupling DFVB with linear solver warm-starts and structural reduction. While warm-starts ensured rapid convergence at each integration step, the model reduction lowered the dimensionality of the optimization problems themselves—and crucially, the most efficient performances reported here were only achievable using the structural insights provided by DFVB. By eliminating hundreds of constraints and variables associated with dynamically inactive basis vectors, the cost per integration step became substantially smaller, and the reduced DFVB approach reached computational speeds highly competitive with standard DFBA. Therefore, DFVB emerges not merely as a dynamic simulator, but as an automated, context-specific pre-processing tool for identifying reduced model topologies.

Beyond numerical efficiency, the simulation-based reduction applied by DFVB proved to be biologically relevant. When compared against *E. coli* transcriptomic profiles, the active reactions identified by DFVB correlated more closely with gene expression than those identified by FVA. The superior F1-score and higher recall suggest that DFVB is highly sensitive to capturing the true representative phenotype under specific environmental conditions. While its aggressive reduction criteria can overestimate the number of inactive reactions, resulting in more false positives than the FVA, the reduced network topology strikes a superior balance for dynamic modeling, retaining essential active pathways while removing irrelevant metabolic functions in the simulation context.

A key virtue of DFVB lies in its capacity for structural regularization during kinetic parameter calibration. Estimating parameters from nonlinear kinetic models in DFBA is notoriously challenging due to solution degeneracy (Sánchez et al. 2014). In the aerobic batch dataset, this was evidenced by the high uncertainty in *K_g_* and *K_ie_* estimation, reflecting a lack of experimental information in the high-inhibition region. Nonetheless, DFVB effectively constrained the kinetic search space and narrowed the confidence intervals. Furthermore, since the experiment was conducted under non-limiting dissolved oxygen conditions, the strict oxygen uptake constrains the saturation of the cellular respiratory capacity (J Nielsen and Villadsen 1994). DFVB captured this bottleneck effectively, forcing the excess carbon flux toward fermentative pathways and resulting in an accurate ethanol production prediction. The virtue of this regularization was even more pronounced in the anaerobic fermentation scenario. DFBA failed to accurately predict ethanol production because its unconstrained LP optimization maximized the growth rate through flux distributions that did not correspond to the actual metabolic phenotype of the yeast. DFVB bypassed this degeneracy by restricting the feasible space to a linear combination of active basis vectors, effectively enforcing a strict physiological coupling between growth and ethanol synthesis. This structural rigidity also enabled a more biologically accurate estimation of the nitrogen yield (*Y_xn_*). It has been demonstrated that yeast biomass yield on nitrogen increases significantly under deficiency due to a drastic reduction in cellular protein content, rising from approximately 19 gDW g^−1^ to 30 gDW g^−1^ (Boer et al. 2003; Varela et al. 2004). The higher *Y_xn_* estimate obtained by DFVB (36.14 gDW g^−1^) successfully captured this adaptive metabolic state. In contrast, DFBA remained trapped in parameter regions associated with nitrogen-rich conditions (14.58 gDW g^−1^), failing to identify the varying yield that governed the culture dynamics. It is important to note, however, that this robust parameter identification relies heavily on the suitability of the initial parameter guess; an incorrect initialization could potentially lock the reduced DFVB model into a biased structural representation.

In summary, DFVB helps bridge the gap between theoretical genome-scale modeling and practical cell culture simulation by providing a mathematically rigorous method to derive low-order, structurally consistent, and computationally efficient dynamic models. Future iterations of the framework could further refine the selection of null-space vectors by incorporating regularization terms into the objective function or by adaptively modifying objectives across different growth phases. Additionally, the reduced-order nature of DFVB facilitates its integration with advanced optimization techniques, such as nonlinear reformulations via KKT conditions, interior-point methods, and hybrid approaches incorporating metabolomic constraints. More importantly, this framework enables the development of more advanced digital models for fermentation optimization and control by allowing robust parameter estimation and more precise predictions.

## Methods

### Flux Balance Analysis

Constraint-based methods are employed for the computation of metabolic fluxes using GEMs (Orth, Thiele, et al. 2010). The vector of fluxes (**v**) is calculated under the steady-state assumption, i.e., there is no significant accumulation of intracellular metabolites. Additionally, reaction fluxes are phenomenologically constrained by upper (**ub**) and lower (**lb**) bounds based on reaction thermodynamics and enzyme capacity:

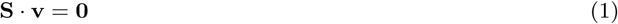

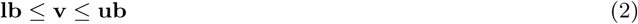

Exploration of the solution space spanned by Eqs. (1–2) can be achieved using optimization. For this task, an objective function that typically maximizes or minimizes a specific flux (e.g., biomass growth) is chosen, leading to the Flux Balance Analysis (FBA) formulation of Eq. (3) (Orth, Thiele, et al. 2010). The objective function choice is not unique, and various studies have proposed alternative objective functions, such as maximizing growth rate and maximizing ATP production, among others (Antoniewicz 2021).

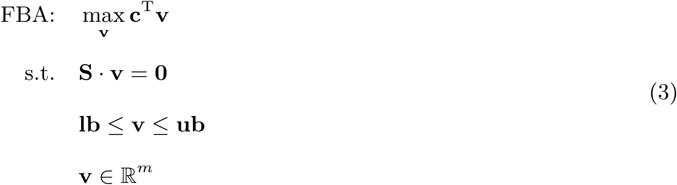

### Dynamic Flux Balance Analysis

A dynamic version of FBA, termed DFBA, was introduced by Mahadevan, Edwards, et al. (2002) that integrates a system of differential equations for the external (unbalanced) species with an FBA problem. Within the static optimization approach of this framework, the FBA solution provides the specific production and consumption rates of the external metabolites at each time step (Eq. 4). These rates are subsequently used to integrate the differential equations and update the bounds of the internal optimization problem.

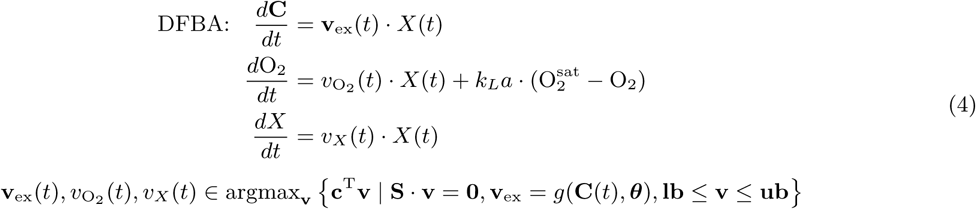

In this framework, **C** represents the vector of extracellular metabolite concentrations, O_2_ the dissolved oxygen concentration, and *X* denotes the biomass concentration. The vector **v**_ex_ corresponds to the subset of exchange fluxes derived from the optimal flux distribution **v**^∗^. This distribution is determined by maximizing a cellular objective subject to the quasi-steady-state mass balance (Eq. 1) and capacity constraints (Eq. 2). The scalars *v*_O₂_ (*t*) and *v_X_* (*t*) represent the oxygen consumption rate and the specific growth rate at time *t*, respectively. Crucially, the system is coupled to the environment through kinetic constraints **v**_ex_ = *g*(**C**(*t*)*, **θ***), where the permissible uptake/secretion rates depend dynamically on the current metabolite concentrations and the particular kinetics encoded in the vector of kinetic parameters ***θ***. Finally, the model incorporates gas-liquid oxygen transfer, defined by the volumetric mass transfer coefficient *k_L_a* and the oxygen saturation concentration 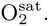.

### Null-space Representation of Mass-balanced Fluxes

The homogeneous linear system **S** · **v** = **0** can be represented using the right null-space of the stoichiometric matrix **S** (Lay et al. 2016). Let **N** be the null-space of **S** and ***α*** be a real-valued vector. Then, a mass-balanced vector **v** lies in the column space of **N** and is given by

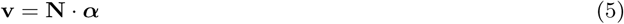

Although the null space of a matrix is a unique linear subspace, it can be spanned by an infinite number of possible bases. To minimize computational cost, a specific basis can be selected based on a sparsity criterion (Saa and LK Nielsen 2016; Bordbar et al. 2014; Gomes de Oliveira Dal’Molin et al. 2018). In this work, the null-space basis of **S** is computed using an algorithm based on Rank-Revealing LU (RRLU) decomposition (Pan 2000). Briefly, the stoichiometric matrix is decomposed as **S** = **LUQ**, where **L** and **Q** are invertible matrices and **U** is an upper triangular matrix that reveals the rank structure. The null-space basis **N** is then constructed from the columns of **Q**^−1^ corresponding to the zero columns of **U**. This approach is selected because RRLU preserves the original matrix entries and structure, thereby facilitating the maintenance of sparsity in the resulting basis and avoiding the computational burden of SVD approaches (Pan 2000).

The null-space basis matrix can be partitioned into rows corresponding to internal reactions (**N**_int_) and external reactions (**N**_ex_). By focusing on the partition of the matrix and the associated fluxes, solutions can be systematically explored using an objective function analogous to that employed in FBA. This approach leads to the formulation of an optimization problem, here termed Flux Vector Balancing (FVB), which is defined as follows:

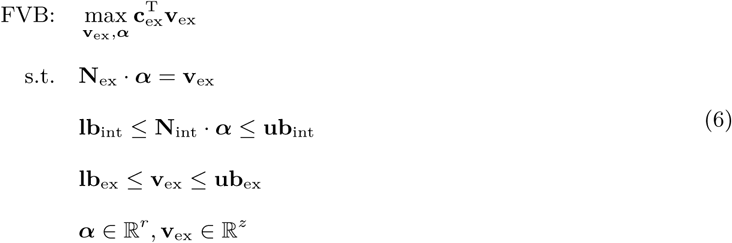

Notably, here the problem is partitioned into internal and external reactions. The vector of coefficients ***α*** carries the weights associated with each basis vector of the null space, describing the activation/inactivation level of each basis vector or flux mode. Simply put, ***α*** operates as a vector of variables that fully parameterize the feasible flux distributions. Internal fluxes are represented as a linear combination of the columns of **N**_int_, whereas the external fluxes are explicitly retained in the formulation to preserve the original structure of the objective function. Here, we have assumed that the optimization target is invariably associated with an external metabolite, as it is common practice for cell culture simulation.

It is crucial to establish that the FVB formulation is not an approximation but a reformulation of the original FBA problem that produces an identical optimal solution (Luenberger and Ye 2008). We provide a rigorous mathematical proof of this proposal in Supplementary File Text S1. Another key insight is the dimensionality reduction of the optimization problem. Let us denote *k* as the rank of **S** and *m* as the total number of reactions, the nullity is given by *m*−*k*. By incorporating *z* exchange reactions explicitly, the FVB problem contains only (*m*−*k*)+*z* variables, compared to *m* in FBA. Given that typically *z* ≪ *m*, this results in a smaller system. Furthermore, the complexity of the constraint set is significantly lower. Standard FBA enforces *n* mass balance equalities and 2*m* bound inequalities, totaling *n* + 2*m* constraints. In contrast, FVB implicitly satisfies the intracellular mass balance, requiring only *z* equality constraints to couple the exchange fluxes (**N**_ex_ · ***α*** = **v**_ex_) alongside the preserved 2*m* bounds. Since the number of exchange reactions is strictly smaller than the number of metabolites (*z < n*), the *x* total constraint count decreases from *n* + 2*m* to *z* + 2*m*. Therefore, this reformulation can effectively leverage the reduced degrees of freedom to improve solver convergence.

### Dynamic Flux Vector Balancing

The Dynamic Flux Vector Balancing (DFVB) algorithm is derived by replacing the internal FBA formulation with the FVB approach (Eq. 7). This modification reduces the size of the internal optimization problem, significantly decreasing the computational demand required to integrate the system of differential equations.

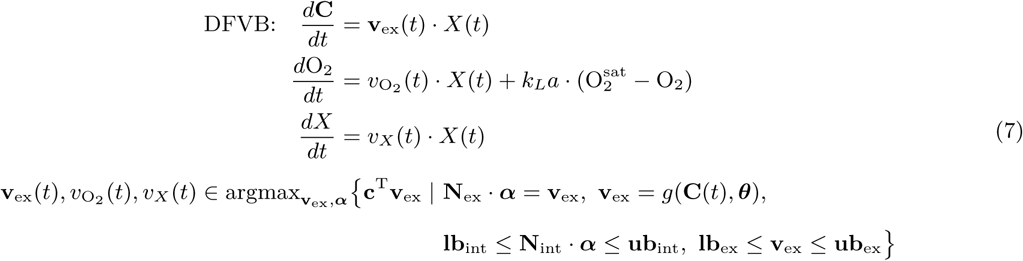

This formulation provides a computationally efficient approach for dynamic metabolic modeling that accounts for oxygen mass transfer, making it particularly suited for large-scale and time-resolved studies in systems biology.

### Simulation-based Model Reduction

DFVB simulations provide time-resolved trajectories for both metabolite concentrations and the flux-basis coefficients ***α***(*t*), under a given scenario captured by the metabolic objective, bounds and kinetic expressions. Since each coefficient *α_j_* (*t*) corresponds to the contribution of the specific null-space vector *j*, its temporal profile directly reflects the activation of the associated flux mode (Gomes de Oliveira Dal’Molin et al. 2018; Bordbar et al. 2014). Basis vectors for which *α_j_* (*t*) = **0** for all *t* ∈ [0*, T*] can therefore be considered inactive throughout the entire simulation.

These inactive basis vectors can be systematically removed from the null-space matrix **N**, yielding a reduced basis that spans only dynamically active metabolic functions. Notably, eliminating columns from **N** may produce rows consisting entirely of zeros; the reactions associated with such rows cannot carry flux under the simulated conditions and are thus also classified as inactive. Once inactive reactions are identified, they can be removed from the original stoichiometric matrix **S**, as well as any metabolite who becomes orphan. This sequential elimination of inactive basis vectors, reactions, and metabolites produces a reduced metabolic model that preserves all dynamically accessible pathways while discarding components that are irrelevant over the entire DFVB trajectory. A schematic overview of the entire DFVB method and model reduction framework is illustrated in Fig. 1. Algorithm 1 formalizes this procedure.

#### Algorithm 1: Model reduction using DFVB-derived activity patterns

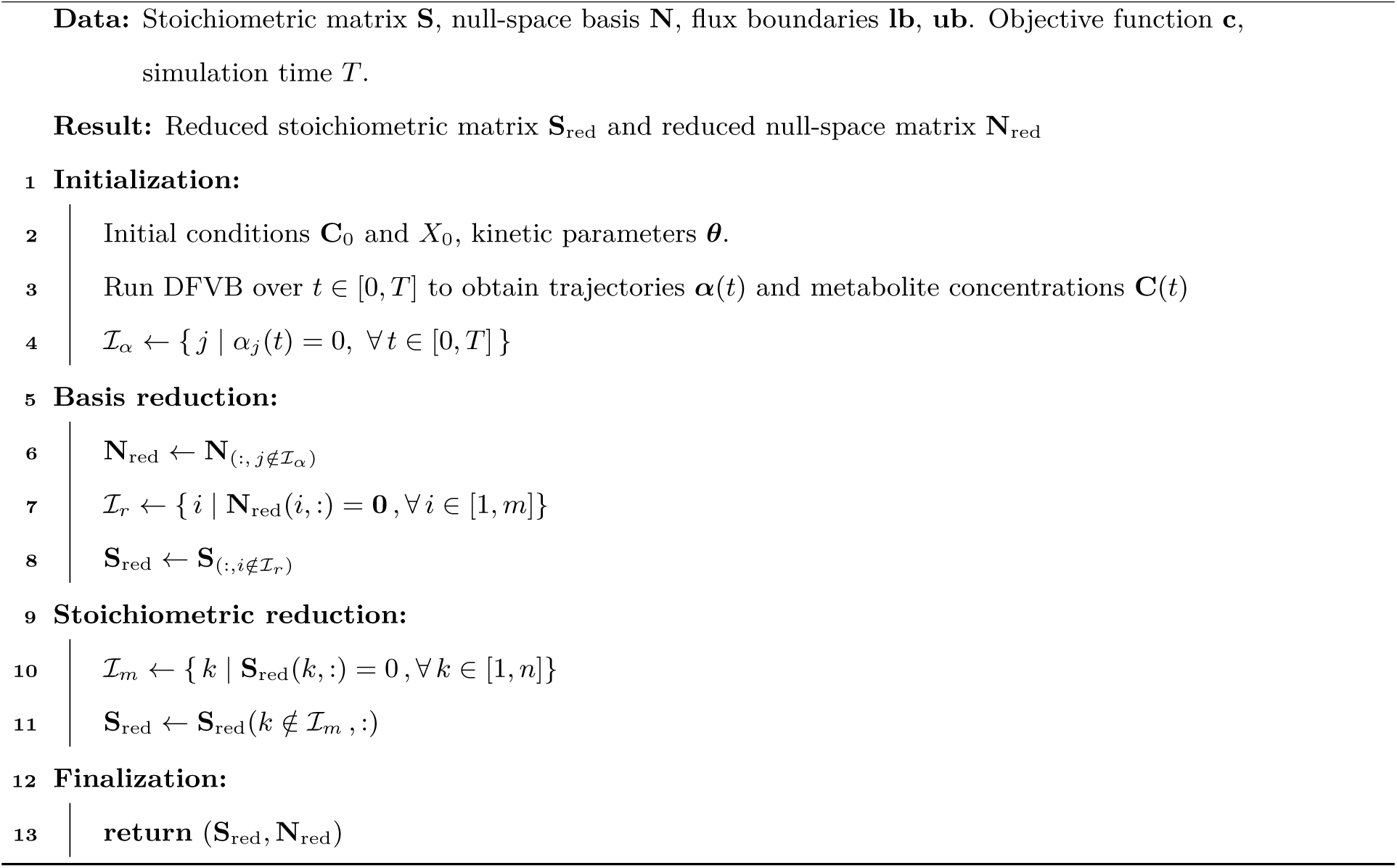

### Evaluation of Dynamic Flux Simulations in Metabolic Models

The computational performance of DFVB was compared against DFBA using metabolic models spanning a wide range of scales. The considered models include the illustrative model from Zhao et al. (2017) (5 metabolites, 7 reactions), which features a single substrate, a product species P, biomass formation, and an internal cycle to explore alternative metabolic routes. To evaluate biologically realistic networks, we employed the *Escherichia coli* core model (73 metabolites, 96 reactions) (Orth, Fleming, et al. 2010), and two genome-scale metabolic models (GEMs): *i*ML1515 for *E. coli* (1878 metabolites, 2713 reactions) (Monk et al. 2017) and Yeast9 for *Saccharomyces cerevisiae* (2806 metabolites, 4131 reactions) (Zhang et al. 2024). The *E. coli* models were modified to include an explicit biomass metabolite and an associated exchange reaction (see Supplementary File Text S2). In all simulations, growth rate maximization was used as the objective function, assuming growth in a minimal medium. Three growth regimes were evaluated: aerobic, anaerobic, and oxygen-limited. Glucose and ammonium were provided as the sole carbon and nitrogen sources, respectively. The bounds for reversible reactions were set to [−1000, 1000] mM gDW^−1^ h^−1^, while irreversible reactions had their lower bound constrained to zero. Oxygen availability was controlled by adjusting the lower bound of the uptake flux. For the aerobic and anaerobic scenarios, this bound was set to −1000 and 0 mM gDW^−1^ h^−1^, respectively. For the oxygen-limited condition, uptake was defined by a Michaelis-Menten expression with a maximum oxygen consumption rate of 20 mM gDW^−1^ h^−1^ and a half-saturation constant of 0.003 mM. In simulations using GEMs, initial conditions were set to 0.5 g L^−1^ for biomass, 1.8 g L^−1^ for glucose, 0.5 g L^−1^ for ammonium, and zero for all metabolic products. Initial dissolved oxygen was set to 0.01 g L^−1^ for the aerobic condition. The volumetric mass transfer coefficient, *k_L_a*, was set to 100 h^−1^, 50 h^−1^, and 0 h^−1^ for the aerobic, oxygen-limited, and anaerobic simulations, respectively, assuming an oxygen saturation concentration of 10 mg L^−1^. Finally, acetate and ethanol uptakes were permitted only if these metabolites were endogenously produced by the metabolic network.

DFVB and DFBA were used to generate dynamic profiles of extracellular metabolites and biomass under different conditions and models. The root-mean-square error (RMSE) was calculated for each metabolite to quantify agreement between DFVB and DFBA and between simulations and experimental data:

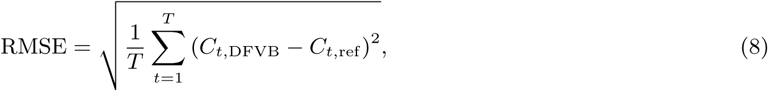

where *C_t,_*_DFVB_ and *C_t,_*_ref_ denote the concentration of a metabolite at time *t* predicted by DFVB and the reference data (either DFBA or experimental measurements), respectively, and *T* is the number of sampled time points.

To accelerate sequential LP solution during dynamic simulations, warm-starts were enabled at every integration step. Providing a feasible basis allows linear solvers to bypass the simplex Phase I procedure and proceed directly to Phase II, substantially reducing the number of pivots required to reach optimality. As reported by Kenefake et al. (2022), warm-starting can substantially reduce computation time while preserving the exact optimal solution, making it particularly effective in DFBA/DFVB simulations where successive LP problems differ only slightly.

Computational performance was evaluated by measuring CPU time for each simulation under four configurations: DFBA and DFVB, with and without solver warm-start, and before and after model reduction. This allowed direct comparison of the impact of both warm-starting and network reduction on the computational cost of dynamic flux simulations.

### Biological Analysis of Model Reduction

To link gene expression levels with the activity of metabolic reactions predicted by simulations using the *i*ML1515 model, the Reaction Activity Score (RAS) was employed (Graudenzi et al. 2018; Huang et al. 2022). This metric assigns an activity value to each reaction based on the expression of its associated genes, accounting for both reactions constrained by the presence of all subunits (logical AND relationships, e.g., multi-unit enzyme complexes) and those involving isozymes (OR relationships).

Transcriptomic data were first normalized using the *rssA* gene as a housekeeping reference, which encodes the 16S rRNA subunit (Peng et al. 2014; Zhou et al. 2011). Subsequently, the Reaction Activity Score (RAS) for each reaction was calculated following the method described by Huang et al. (2022). Formally, letting *I* denote the set of genes encoding enzymes associated with metabolic reactions, the expression level of each gene was initially normalized according to the number of reactions in which the corresponding enzyme participates, yielding mRNA_norm_*_,j_*. The specific score *RAS_j_* was then computed depending on the catalytic requirements of the reaction: for those catalyzed by a single enzyme, the score was defined directly as *RAS_j_* = mRNA_norm_*_,j_* ; for reactions involving an OR relationship among the genes in set *I* (e.g., isozymes), it was calculated as the sum of the normalized expressions, 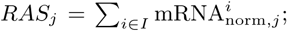 whereas for reactions involving an AND relationship (e.g., multi-subunit enzyme complexes), the score was determined by the minimum expression level among the required subunits, 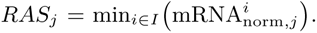 Finally, reactions lacking associated genes were excluded from the analysis, as their activity could not be reliably verified in the absence of a corresponding gene-protein-reaction mapping. After computing the RAS, reactions were classified as active/inactive using a dynamic threshold at the 40th percentile, also corresponding to a bimodal separation (see Supplementary File Text S3). This cutoff preserves essential metabolic functionalities often lost with stricter thresholds, while effectively filtering transcriptional noise (Opdam et al. 2017).

We employed a curated subset of the PRECISE compendium generated by Sastry et al. (2019). While the full dataset contains 278 RNA-seq profiles spanning diverse environmental and regulatory conditions, 49 transcriptomes were selected, corresponding specifically to *Escherichia coli* K-12 MG1655 grown aerobically in M9 minimal medium with glucose and ammonium as the sole carbon and nitrogen sources. In the original study, samples were harvested during exponential growth (OD_600_ ≈ 0.5), subjected to standardized RNA extraction and rRNA depletion protocols, and sequenced using the Illumina platform. After quality-controlled processing and log-transformation of transcript abundances, the resulting dataset delivers high-quality expression measurements for 1516 enzyme-associated genes. This controlled and internally consistent subset was chosen to rigorously evaluate whether the reduced metabolic models preserve functional coherence at the gene-reaction level.

### Kinetic Parameter Estimation for Modeling Culture Dynamics

To demonstrate the practical advantages of the DFVB framework over conventional DFBA, kinetic parameter calibration was performed in two distinct fermentation scenarios. In standard DFBA, estimating kinetic parameters from experimental data is notoriously challenging and computationally expensive (Sánchez et al. 2014). By structurally reducing the metabolic network and isolating the dynamically active subspace, DFVB significantly decreases this computational burden, enabling a robust and more efficient parameter estimation process.

The kinetic models calibrated for the aerobic batch culture of *S. cerevisiae* and *Sauvignon Blanc* fermentation are based on Sánchez et al. (2014) and Zenteno et al. (2010), respectively, and they are described in detail in Supplementary File Text S4.

For the aerobic batch fermentation of *S. cerevisiae*, the calibrated parameters included the maximum glucose uptake rate (*v_S,_*_max_), the half-saturation constant for glucose (*K_S_*), the ethanol inhibition constant (*K_ie_*), the lower bound on oxygen uptake in the metabolic model (*lb*_O₂_), and the biomass yield coefficients for glycerol, citrate, and lactate (*Y*_x,gly_, *Y*_x,cit_, *Y*_x,lac_).

For the *Sauvignon Blanc* fermentation model, the calibrated parameters were the maximum specific growth rate (*µ*_0_); the half-saturation constants for nitrogen, glucose, and fructose uptake (*K_n,_*_0_, *K_g,_*_0_, *K_f,_*_0_); the ethanol inhibition constant (*K_ie,_*_0_); the biomass yields on nitrogen, glucose, and fructose (*Y_xn_*, *Y_xg_*, *Y_xf_*); the ethanol yield on fructose (*Y_ef_*); and the baseline production rates of ethanol from glucose and fructose (*β_g,_*_0_, *β_f,_*_0_).

The parameter calibration was formulated as an iterative optimization problem. Here, the decision variable vector ***θ*** represents the kinetic parameters governing the system dynamics (Eq. 4 and Eq. 7). Each optimization step involved executing a forward simulation using either the DFBA or DFVB framework to minimize a loss function based on the relative mean squared error between the predicted and experimentally measured concentrations. A detailed mathematical description of this optimization problem is provided in Supplementary File Text S5. For both kinetic models, the optimization was initialized using parameter estimates from the literature (see Supplementary File Text S4). The initial conditions were also included as fitted parameters, as experimental error in these measurements can severely affect parameter calibration (Cao et al. 2011). Due to the lack of replicates in the Sánchez et al. (2014) dataset, a variation of 30% relative to the reported values was allowed to constrain the estimation. In contrast, for the *Sauvignon Blanc* parameter calibration, the bounds were defined using two standard deviations derived from the three experimental replicates, thereby explicitly accounting for measurement uncertainty.

Finally, a bootstrap analysis with 1000 resampling iterations was performed for both calibration procedures (Wong et al. 2016). In each iteration, the experimental dataset was resampled with replacement, and the model was recalibrated using the resampled data. The resulting parameter distributions were then used to compute the mean estimates associated 95% bootstrap confidence intervals, enabling a robust comparison of the parameter uncertainty and overall calibration performance of DFBA and DFVB.

### Experimental Methods

Two independent experimental datasets were used to evaluate parameter fitting of dynamic models. The first dataset consisted of laboratory-scale fermentations of *Sauvignon Blanc* must (blend-quality lot sourced from Fundo San Ignacio, Lontué, Chile), carried out in 2-L jacketed bioreactors in triplicate. The system followed a stepwise temperature profile of 19, 16, and 21 ^◦^C, regulated by a circulating water bath connected to the vessel jackets, with temperature logging but no closed-loop control. Prior to filling, the must was homogenized, supplemented with 50 mg L^−1^ SO_2_, as potassium metabisulfite, and adjusted to a pH of 3.45 using tartaric acid. Fermentations were inoculated with *Saccharomyces cerevisiae* VL3 (Laffort, France). Nitrogen supplementation consisted of two aqueous bolus additions of diammonium phosphate (10% w/w stock), applied at inoculation and mid-fermentation. The reactors were magnetically stirred at 400 rpm for 2 min and samples were collected one to two times per day via the top port. Biomass concentration and viability were quantified from unprocessed aliquots using the Oculyze system with 0.01% alkaline methylene violet, while dry cell weight was determined gravimetrically after drying cell pellets at 60 ^◦^C for 48 h. All other analytes were measured from the supernatant obtained by centrifugation (5 min, 4200 rpm). These measurements included density (Anton Paar DM35), ethanol (Anton Paar Alcolyzer), and enzymatic assays (Biosystems Y15) for glucose, fructose, yeast-assimilable nitrogen, glycerol, and organic acids. The progression of these fermentations to complete dryness (*<*2 g L^−1^ residual sugars) provided a high-resolution experimental dataset. By capturing the complex transient dynamics of anaerobic growth under shifting temperature regimes, this experimental setup provides sufficient data to benchmark the parameter identification capabilities of both DFBA and DFVB frameworks in highly non-linear scenarios.

The second dataset was taken from Sánchez et al. (2014), who reported controlled aerobic batch fermentations of *Saccharomyces cerevisiae* using the EC1118 wine yeast strain (Lalvin, Switzerland). The cultures were initiated with 20 g L^−1^ glucose in defined minimal medium and performed in 1-L stirred-tank bioreactors equipped with temperature, pH, and dissolved oxygen control. Samples collected over 12 h enabled quantification of key extracellular metabolites, including glucose, ethanol, glycerol, citrate, and lactate through high-performance liquid chromatography (HPLC). This dataset provides high-resolution time-course measurements of biomass and metabolite dynamics under aerobic condition, serving as an independent source for evaluating parameter identifiability and model predictive capability.

### Computational Implementation

The algorithms were implemented and executed in MATLAB 2025a (The MathWorks, Natick, MA, USA). Linear programming problems were solved using Gurobi Optimizer 12.0.0 (Gurobi Optimization, Inc., Houston, TX, USA) with both relative and absolute tolerances set to 10^−8^. To enable the warm-start configuration in Gurobi, the optimal basis from the previous LP (vbasis and cbasis) is loaded, and the LPWarmStart parameter is set to 1. Ordinary differential equations were integrated using MATLAB’s built-in *ode113* solver, a variable-step, variable-order Adams–Bashforth–Moulton method (Shampine and Reichelt 1997). Parameter estimation was carried out using the local optimizer *fminsearchbnd* (D’Errico 2026), a bound-constrained extension of the standard Nelder-Mead simplex algorithm implemented in MATLAB. This derivative-free method circumvents the limitations of unconstrained optimization by applying internal coordinate transformations to the decision variables. This approach enables rigorous enforcement of lower and upper bounds on the kinetic parameters without computing gradients, ensuring that the calibrated values remain within biologically realistic ranges throughout the optimization process. Simulations were carried out on a laptop equipped with an Intel Core Ultra i7 155H processor, 16 GB RAM, 64-bit architecture, running Windows 11. Bootstrapping calibrations were run on a node of the Unix-based Cluster of the School of Engineering at the Pontificia Universidad Católica de Chile (more details for the computational resources employed can be found in Supplementary File Text S6). The code and model reconstructions used in this study are available in the GitHub repository (https://github.com/SysBioengLab/DFVB).

## Data access

All data accompanying this research are presented directly in the manuscript and Supplementary Material. The complete datasets, source code, and implementation details required to reproduce this work are publicly available in the GitHub repository: https://github.com/SysBioengLab/DFVB.

## Acknowledgments

PAS and JRP acknowledge the financial support from ANID – Applied Research Center – CIA 250013. IT and PAS also acknowledge the support from the UC School of Engineering through the Seed Fund ILO 2025.

## Author Contributions

**Ignacio Tapia**: Conceptualization, Investigation, Methodology, Formal analysis, Software, Writing – original draft, Writing – review & editing. **Cristóbal Torrealba**: Investigation, Methodology, Validation, Writing-review & editing. **Ricardo Luna**: Funding, Writing–review & editing. **José Ricardo Pérez**: Conceptualization, Investigation, Methodology, Formal analysis, Writing – original draft, Writing – review & editing. **Pedro Saa**: Conceptualization, Investigation, Methodology, Formal analysis, Writing – original draft, Writing – review & editing.

## References

Altamirano Á, I Tapia, V Acuña, D Garrido, and PA Saa. 2026. METACONE: A scalable framework for exploring the conversion cone of metabolic networks. Computational Biology and Chemistry 120:108607.

Antoniewicz MR. 2021. A guide to metabolic flux analysis in metabolic engineering: Methods, tools and applications. Metabolic Engineering 63:2–12.

Ataman M, DF Hernandez Gardiol, G Fengos, and V Hatzimanikatis. 2017. redGEM: Systematic reduction and analysis of genome-scale metabolic reconstructions for development of consistent core metabolic models. PLOS Computational Biology 13:1–22.

Baroukh C, R Muñoz-Tamayo, J.-P Steyer, and O Bernard. 2014. DRUM: A New Framework for Metabolic Modeling under Non-Balanced Growth. Application to the Carbon Metabolism of Unicellular Microalgae. PLOS ONE 9:1–15.

Boer VM, JH de Winde, JT Pronk, and MD Piper. 2003. The genome-wide transcriptional responses of *Saccharomyces cerevisiae* grown on glucose in aerobic chemostat cultures limited for carbon, nitrogen, phosphorus, or sulfur. The Journal of Biological Chemistry 278:3265–3274.

Bordbar A, H Nagarajan, NE Lewis, H Latif, A Ebrahim, S Federowicz, J Schellenberger, and BO Palsson. 2014. Minimal metabolic pathway structure is consistent with associated biomolecular interactions. Molecular Systems Biology 10:737.

Cao J, L Wang, and J Xu. 2011. Robust Estimation for Ordinary Differential Equation Models. Biometrics 67:1305–1313.

D’Errico J. 2026. fminsearchbnd, fminsearchcon. MATLAB Central File Exchange.

de Oliveira RD, GA Le Roux, and R Mahadevan. 2023. Nonlinear programming reformulation of dynamic flux balance analysis models. Computers & Chemical Engineering 170:108101.

Dromms RA, JY Lee, and MP Styczynski. 2020. LK-DFBA: a linear programming-based modeling strategy for capturing dynamics and metabolite-dependent regulation in metabolism. BMC Bioinformatics 21:93.

Erdrich P, R Steuer, and S Klamt. 2015. An algorithm for the reduction of genome-scale metabolic network models to meaningful core models. BMC Systems Biology 9:48.

Gomes de Oliveira Dal’Molin C, L.-E Quek, PA Saa, R Palfreyman, and LK Nielsen. 2018. From reconstruction to C4 metabolic engineering: A case study for overproduction of polyhydroxybutyrate in bioenergy grasses. Plant Science 273:50–60.

Gotsmy M, D Giannari, R Mahadevan, and J Zanghellini. 2024. Optimizing Fed-Batch Processes with Dynamic Control Flux Balance Analysis. IFAC-PapersOnLine 58:109–114.

Graudenzi A, D Maspero, M Di Filippo, M Gnugnoli, C Isella, G Mauri, E Medico, M Antoniotti, and C Damiani. 2018. Integration of transcriptomic data and metabolic networks in cancer samples reveals highly significant prognostic power. Journal of Biomedical Informatics 87:37–49.

Gu C, GB Kim, WJ Kim, HU Kim, and SY Lee. 2019. Current status and applications of genome-scale metabolic models. Genome Biology 20:121.

Höffner K, SM Harwood, and PI Barton. 2013. A reliable simulator for dynamic flux balance analysis. Biotechnology and Bioengineering 110:792–802.

Huang Y, V Mohanty, M Dede, M Daher, L Li, K Rezvani, and K Chen. 2022. Characterizing metabolism from bulk and single-cell RNA-seq data using METAFlux. bioRxiv doi: 10.1101/2022.05.18.492580.

Kenefake D, E Armingol, NE Lewis, and EN Pistikopoulos. 2022. An improved algorithm for flux variability analysis. BMC bioinformatics 23:550.

Lay D, S Lay, and J McDonald. 2016. Linear Algebra and Its Applications. Pearson.

Luenberger DG and Y Ye. Linear and Nonlinear Programming. In: 3rd ed. International Series in Operations Research & Management Science. Chap. 6.5, pp. 179–181.

Mahadevan R and C Schilling. 2003. The effects of alternate optimal solutions in constraint-based genome-scale metabolic models. Metabolic Engineering 5:264–276.

Mahadevan R, JS Edwards, and FJD III. 2002. Dynamic flux balance analysis of diauxic growth in Escherichia coli. Biophysical Journal 83:1331–1340.

Martínez VS, PA Saa, J Jooste, K Tiwari, L.-E Quek, and LK Nielsen. 2022. The topology of genome-scale metabolic reconstructions unravels independent modules and high network flexibility. PLOS Computational Biology 18:1–20.

Monk JM et al. 2017. iML1515, a knowledgebase that computes Escherichia coli traits. Nature biotechnology 35:904–908.

Nielsen J and J Villadsen. 1994. Bioreaction Engineering Principles. Springer New York, New York, NY.

Opdam S, A Richelle, B Kellman, S Li, DC Zielinski, and NE Lewis. 2017. A systematic evaluation of methods for tailoring genome-scale metabolic models. Cell systems 4:318–329.

Orth J, R Fleming, and B Palsson. 2010. Reconstruction and Use of Microbial Metabolic Networks: the Core *Escherichia coli* Metabolic Model as an Educational Guide. EcoSal Plus 4:10.1128/ecosalplus.10.2.1.

Orth J, I Thiele, and B Palsson. 2010. What is flux balance analysis. Nature biotechnology 28:245–248.

Pan CT. 2000. On the existence and computation of rank-revealing LU factorizations. Linear Algebra and its Applications 316:199–222.

Peng S, R Stephan, J Hummerjohann, and T Tasara. 2014. Evaluation of three reference genes of Escherichia coli for mRNA expression level normalization in view of salt and organic acid stress exposure in food. FEMS Microbiology Letters 355:78–82.

Pinhal S, D Ropers, J Geiselmann, and H de Jong. 2019. Acetate Metabolism and the Inhibition of Bacterial Growth by Acetate. Journal of Bacteriology 201:10.1128/jb.00147–19.

Saa PA and LK Nielsen. 2016. Fast-SNP: a fast matrix pre-processing algorithm for efficient loopless flux optimization of metabolic models. Bioinformatics 32:3807–3814.

Sánchez BJ, JR Pérez-Correa, and E Agosin. 2014. Construction of robust dynamic genome-scale metabolic model structures of Saccharomyces cerevisiae through iterative re-parameterization. Metabolic Engineering 25:159–173.

Sastry AV, Y Gao, R Szubin, Y Hefner, S Xu, D Kim, KS Choudhary, L Yang, ZA King, and BO Palsson. 2019. The Escherichia coli transcriptome mostly consists of independently regulated modules. Nature Communications 10:5536.

Schroeder W and R Saha. 2020. Introducing an Optimization- and explicit Runge-Kutta- based Approach to Perform Dynamic Flux Balance Analysis. Scientific Reports 10:9241.

Scott F, P Wilson, R Conejeros, and VS Vassiliadis. 2018. Simulation and optimization of dynamic flux balance analysis models using an interior point method reformulation. Computers & Chemical Engineering 119:152–170.

Shampine LF and MW Reichelt. 1997. The MATLAB ODE Suite. SIAM Journal on Scientific Computing 18:1–22.

Singh D and MJ Lercher. 2020. Network reduction methods for genome-scale metabolic models. Cellular and Molecular Life Sciences 77:481–488.

Smallbone K and E Simeonidis. 2009. Flux balance analysis: A geometric perspective. Journal of Theoretical Biology 258:311–315.

Varela C, F Pizarro, and E Agosin. 2004. Biomass Content Governs Fermentation Rate in Nitrogen-Deficient Wine Musts. Applied and Environmental Microbiology 70:3392–3400.

Wiback SJ, R Mahadevan, and BØ Palsson. 2003. Reconstructing metabolic flux vectors from extreme pathways: defining the *α*-spectrum. Journal of Theoretical Biology 224:313–324.

Wong RKW, CB Storlie, and TCM Lee. 2016. A Frequentist Approach to Computer Model Calibration. Journal of the Royal Statistical Society Series B: Statistical Methodology 79:635–648.

Zenteno MI, JR Pérez-Correa, CA Gelmi, and E Agosin. 2010. Modeling temperature gradients in wine fermentation tanks. Journal of Food Engineering 99:40–48.

Zhang C et al. 2024. Yeast9: a consensus genome-scale metabolic model for S. cerevisiae curated by the community. Molecular Systems Biology 20:1134–1150.

Zhao X, S Noack, W Wiechert, and EV Lieres. 2017. Dynamic flux balance analysis with nonlinear objective function. Journal of mathematical biology 75:1487–1515.

Zhou K, L Zhou, J Lim, R Zou, G Stephanopoulos, and H.-P Too. 2011. Novel reference genes for quantifying transcriptional responses of Escherichia coli to protein overexpression by quantitative PCR. BMC molecular biology 12:18.

